# Scene context and attention independently facilitate MEG decoding of object category

**DOI:** 10.1101/2024.06.30.601374

**Authors:** Olga Leticevscaia, Talia Brandman, Marius V. Peelen

## Abstract

Many of the objects we encounter in our everyday environments would be hard to recognize without any expectations about these objects. For example, a distant silhouette may be perceived as a car because we expect objects of that size, positioned on a road, to be cars. Reflecting the influence of such expectations on visual processing, neuroimaging studies have shown that when objects are poorly visible, expectations derived from scene context facilitate the representations of these objects in visual cortex from around 300 ms after scene onset. The current magnetoencephalography (MEG) study tested whether this facilitation occurs independently of attention and task relevance. Participants viewed degraded objects alone or within their original scene context while they either attended the scenes (attended condition) or the fixation cross (unattended condition), temporally directing attention away from the scenes. Results showed that at 300 ms after stimulus onset, multivariate classifiers trained to distinguish clearly visible animate vs inanimate objects generalized to distinguish degraded objects in scenes better than degraded objects alone, despite the added clutter of the scene background. Attention also modulated object representations at this latency, with better category decoding in the attended than the unattended condition. The modulatory effects of context and attention were independent of each other. Finally, data from the current study and a previous study were combined (N=51) to provide a more detailed temporal characterization of contextual facilitation. These results extend previous work by showing that facilitatory scene-object interactions are independent of the specific task performed on the visual input.

## Introduction

Scene context can have a powerful influence on object perception. For example, objects presented within a coherent scene context (such as a car on a road) are recognized more easily than objects presented within an incoherent scene context (such as a car in a kitchen; Bar, 2004; Biederman et al., 1982; Davenport & Potter, 2004; Oliva & Torralba, 2007). Scene context also guides attention to objects (Castelhano & Krzys, 2020; Wu et al., 2014). Multiple types of contextual factors have been distinguished and shown to independently contribute to object recognition and attentional guidance, including the probability that an object would occur in a particular scene and the typical position and size of an object within a scene (Biederman et al., 1982; Castelhano & Krzys, 2020; Wu et al., 2014). Finally, when objects are poorly visible (such as when they are far away, in the periphery, or partially occluded) or viewed only briefly, expectations derived from scene context make objects look less blurry (Rossel et al., 2022, 2023).

In line with the finding of perceptual sharpening (Rossel et al., 2022, 2023), neuroimaging studies have shown that expectations derived from scene context facilitate object representations in visual cortex (Peelen et al., 2024). For example, when a poorly visible object is presented within its original scene context, multivariate functional magnetic resonance imaging (fMRI) activity patterns in object-selective cortex become more similar to activity patterns evoked by the intact version of that object, as compared to when the object is presented without scene context (Brandman & Peelen, 2017). Using the same experimental approach, scene-based facilitation of object representations was also found using multivariate analysis of magnetoencephalography (MEG) data. Results showed that the contextual facilitation effect emerged relatively late, from around 320 ms after stimulus onset, likely reflecting top-down feedback to visual cortex (Brandman & Peelen, 2017).

Here, we follow up on this work, asking whether the scene-based facilitation of object representations, observed at 320 ms after stimulus onset, depends on attention and the task relevance of these objects. In our previous work, participants paid full attention to the objects, both in space and in time, to discriminate objects from numbers (Brandman & Peelen, 2017). Spatially, images were always presented at the same central location and participants distributed their attention across the image. Temporally, images were presented at a fixed delay relative to the onset of a fixation cross so that their onset was fully predictable. This ensured that participants were optimally prepared to process the scenes and the objects within them. However, in daily life, most objects in scenes are not directly relevant to ongoing task performance, and often objects appear at locations and moments that are not fully known in advance (such as a car suddenly appearing from a driveway). This raises the question of whether scene-based expectations modulate object processing also when objects are not directly task relevant and when, accordingly, attention is not fully focused on these objects.

Previous behavioral and EEG studies investigating the impact of semantic scene-object congruency (e.g., a car on a road vs a car in a kitchen) have shown that these congruency effects may be relatively independent of attention and task relevance (but see Gronau, 2021). For example, in a behavioral study, congruent (vs incongruent) scene context facilitated object naming independently of whether spatial attention was focused on the foreground object or the background scene (Munneke et al., 2013). Another study showed that objects that violate scene-based expectations (e.g., a toothbrush in an office) attracted fixations even when the scenes (and objects) were entirely irrelevant for the task (Cornelissen & Võ, 2017). In line with these findings, event-related potential (ERP) components indexing semantic scene-object violations (N300/400; Võ & Wolfe, 2013) were found both when objects were task relevant and when they were task-irrelevant (Chen et al., 2022).

While these studies demonstrate that semantic violations are noticed relatively automatically, they do not speak to the question of whether scene-based predictions facilitate object representations independently of attention and task relevance. These two processes – the detection of semantic violations and the modulation of object representations – likely reflect different neural mechanisms (Chen et al., 2022). For example, in a recent EEG study (Chen et al., 2022), N300/400 responses to semantic violations were observed irrespective of task relevance but object representations (indexed by a decoding analysis) were only facilitated by scene context for objects that were task-relevant. Based on these results, the authors concluded that scene-based predictions about object properties are gated by task demands, such that predictions are only generated for objects for which the perception of object detail is task-relevant (Chen et al., 2022).

Notably, however, previous EEG studies all investigated the influence of scene context on the processing of clearly visible (i.e., unambiguous) objects. This is important because context-based expectations may facilitate object processing only when object features need to be disambiguated, in line with predictive processing accounts of perception (Peelen et al., 2024). For example, the effect of scene context on the perceived sharpness of objects in a behavioral study was shown to depend on the reliability of the visual input: when visual input was unreliable (e.g., a blurred object), scene context increased its perceived sharpness but when visual input was reliable this effect reversed in favor of scene-inconsistent objects (Rossel et al., 2023). Similarly, in a challenging object categorization task, scene context facilitated the categorization of blurry objects more than it facilitated the categorization of intact objects (Brandman & Peelen, 2017). Therefore, it remains to be investigated whether scene-based facilitation of ambiguous visual object representations is independent of attention and task relevance.

Here, we adapted our MEG paradigm (Brandman & Peelen, 2017) to include an attention manipulation to address this question. In different experimental runs, participants either attended the scene/object images (*attended condition*), pressing a button when a number appeared instead of an image, or attended the fixation cross (*unattended condition*), pressing a button in response to a small luminance change. To further draw attention away from image processing in the unattended condition, the luminance change always occurred just before image onset, thereby manipulating temporal attention (Nobre & Van Ede, 2018). This way, in the unattended condition, the scene/object image was task-irrelevant, with attention spatially focused on the fixation cross (rather than the whole image) and temporally directed away from image onset.

Closely following the analysis approach of our previous MEG study, we used time-sensitive cross-decoding analyses in which we classified the category of the objects using a classifier trained on the multivariate MEG response patterns evoked by intact objects. This classifier was then tested on response patterns evoked by degraded objects presented alone or in scenes, while participants either attended the image or the fixation cross (Figure 1A). We hypothesized that object decoding would be better for objects in scenes than for objects alone at around 320 ms after image onset, replicating our previous finding (Brandman & Peelen, 2017). Furthermore, in line with the behavioral findings reviewed above ; Cornelissen & Võ, 2017), we hypothesized that this modulation would be independent of attention. Finally, we hypothesized that attention would also modulate object decoding, with better decoding of attended than unattended objects (Kaiser et al., 2016; Mangun & Hillyard, 1991; Rohenkohl et al., 2012).

**Figure 1.**
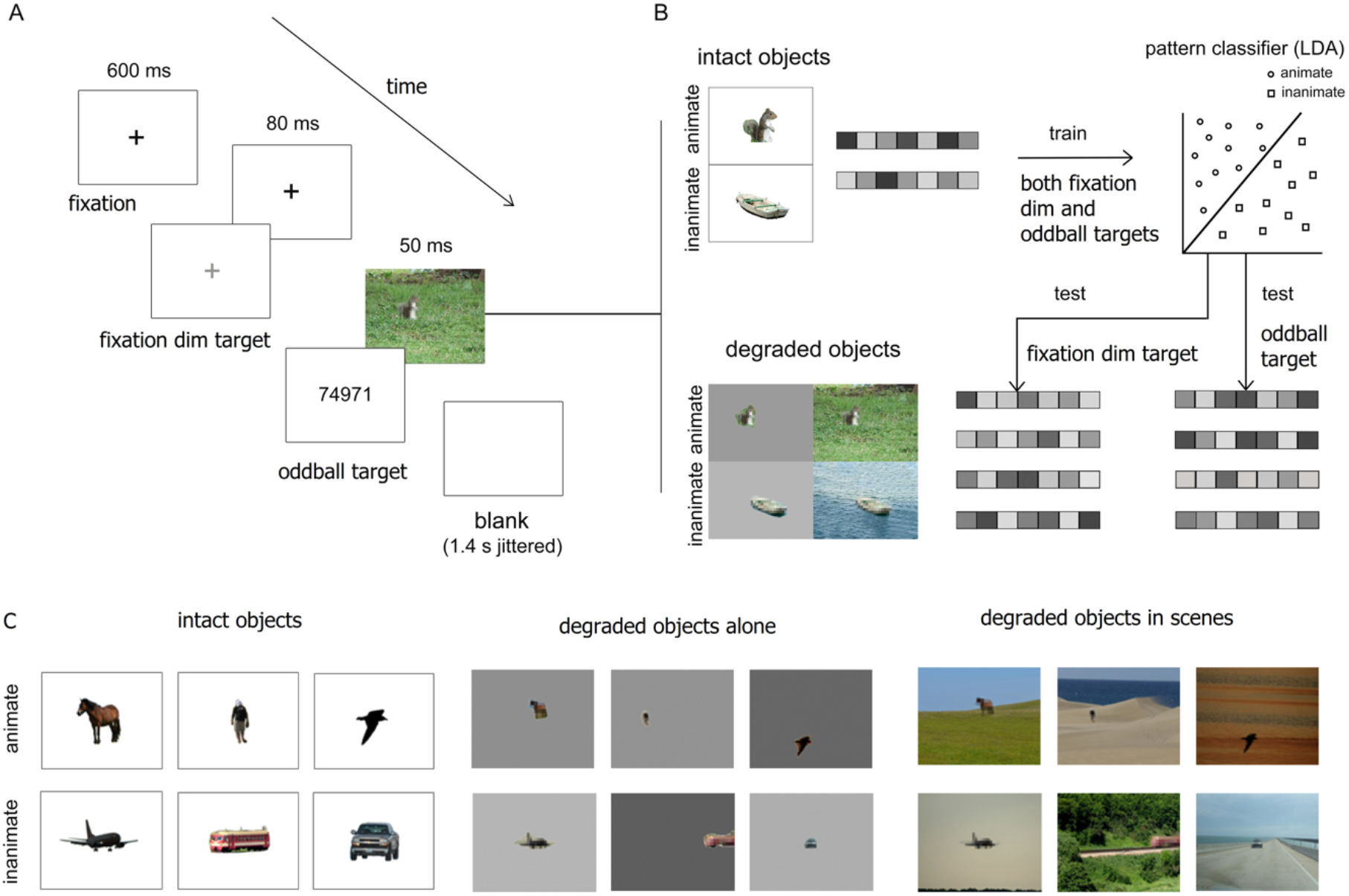
Illustration of experimental design and data analysis. A. Example of trial sequence. Participants viewed a 600 ms black fixation cross followed by an 80 ms fixation cross that was either the same or slightly dimmed. Fixation was followed either by an object or an oddball target (random number) for 50 ms. In the attended condition, participants were instructed to attend the object images and press a button when the oddball target appeared instead of an image. In the unattended condition, participants were instructed to attend the fixation cross and press a button when it dimmed, thereby temporally directing attention away from image onset; B. Cross-decoding analysis. An LDA classifier was trained to discriminate between animate and inanimate intact objects based on the response patterns across MEG sensors, and tested on animacy discrimination of degraded objects, separately for each of the four main experiment conditions, i.e., either with or without scene background, in either the temporally attended (detect oddball targets) or unattended (detect fixation dim targets) condition; C. Examples of images in the different conditions. The main experiment contained 60 unique images (see also Brandman & Peelen, 2017).

## Materials and Methods

### Participants

Twenty-six healthy participants (14 female, mean 25.6 years ± 3.39 SD) were included in this study. All participants had normal or corrected-to-normal vision and gave informed consent. Sample size was matched to that of our previous MEG study (Brandman & Peelen, 2017). Three additional participants took part but were excluded from data analysis, one because of low performance in the unattended condition (<20% correct), one due to excessive movement and signal noise during online inspection, and one due to an online technical issue, which prevented the completion of data acquisition. All procedures were approved by the ethics committee of the University of Trento.

### Stimuli

The stimulus set for the degraded object experiment consisted of degraded objects that were perceived as ambiguous on their own but that were easily categorized when presented in context (Brandman & Peelen, 2017). This set included photographs of 30 animate (animals and people) and 30 inanimate (cars, boats, planes, and trains) objects in outdoor scenes. Each photograph included a single foreground object, which was blurred relative to the scene (which was not blurred). Each object appeared in two images: once in its original background, and once on a uniform gray background of mean luminance of the original background, resulting in a total of 120 images. Objects were presented in their original location both when presented alone and in a scene. To avoid familiarity effects passing from objects in scenes to objects alone, the stimulus set was split in two, such that different objects were presented for degraded objects in scenes and for degraded objects alone within a given subject. (For example, for two stimuli Bird1 and Bird2, a given participant would either view Bird1 in isolation and Bird2 in the sky, or vice versa). The two sets were counterbalanced across subjects.

The stimulus set for the intact object experiment (used for classifier training; see also Brandman & Peelen, 2017) included the 60 objects from the main experiment and an additional set of 60 new objects that were matched for category and subcategory of the main experiment set, all in high resolution and centred on a white background. All stimuli were presented at 5.99° × 4.50° visual degrees (400 × 300 pixels).

### Experimental design

In each trial, participants viewed a black fixation cross (600 ms), which was either followed by a gray (RGB [230, 230, 230]) fixation cross or remained black (80 ms), then followed by an object image (50 ms) or an oddball target (a randomly generated number of five digits), and a mean inter-stimulus interval of 1.4s ±500 ms jitter (Figure 1A).

Attention to objects was manipulated by diverting participants’ attention to different parts of a trial in different tasks; trial sequence was identical in both tasks. To draw attention to objects (*attended condition*), participants were asked to attend object images and press a button when they saw a number, which randomly appeared instead of an object on average once every 16 trials (oddball task). To divert participants’ attention temporally away from objects (*unattended condition*), participants were instructed to attend the fixation cross which appeared prior to object images, and to press a button when they detected a change in its luminance (fixation-dimming task; adapted from Stein & Peelen, 2017). The two tasks alternated across runs.

The degraded object experiment consisted of 8 runs (4 runs of attended and 4 runs of unattended condition) of 400 s duration, each composed of 7 fixation breaks (8 s fixation screen each), 8 oddball trials, and 30 trials per condition: animate / inanimate × object alone / object-in-scene (128 trials/run). The fixation cross dimmed in 24 trials per run. The intact object experiment consisted of 4 runs (2 runs of attended and unattended conditions each) of 400 s duration, each composed of 7 fixation breaks (8 s each), 8 oddball trials, and 60 trials per condition: animate / inanimate. The fixation cross dimmed in 24 trials per run. The composition of each run was the same in the attended and unattended conditions, differing only in task instructions.

Prior to the experiment, all participants completed a practice session of the degraded object experiment to ensure they understood both tasks. The practice was performed once while participants responded to oddball images in the attended condition, and once while participants responded to dimmed fixations in the unattended condition. Each condition included 16-32 trials.

### Data acquisition and pre-processing

Electromagnetic brain activity was recorded using an Elekta Neuromag 306 MEG system, composed of 204 planar gradiometers and 102 magnetometers. Signals were sampled continuously at 1000 Hz and bandpass filtered online between 0.1 and 330 Hz. Offline preprocessing was done using the Elekta MaxFilter/MaxMove software, MATLAB (RRID:SCR_001622) and the FieldTrip analysis package (RRID:SCR_004849). Signals were filtered of external noise and spatially realigned with the Maxwell Filter and Move, using signal space separation algorithms (Taulu & Simola, 2006). Artifact removal was performed manually by excluding noisy trials and channels. Data were then demeaned, detrended, down-sampled to 100 Hz, and time-locked to stimulus onset. The data were averaged across trials of the same exemplar across runs, resulting in a total of 120 unique test trials throughout the degraded object experiment (30 per condition), and 120 unique train/test trials throughout the intact object experiment (30 per condition).

### Multivariate analysis

Multivariate analysis was performed using the CoSMoMVPA toolbox (Oosterhof et al., 2016; RRID:SCR_014519). Following Brandman and Peelen (2017), decoding was performed across posterior magnetometers (48 channels) of each participant, between 0 and 500 ms. Prior to decoding, temporal smoothing was applied by averaging across neighbouring time points with a radius of 2 (20 ms). A linear discriminant analysis (LDA) classifier was trained to discriminate between response patterns to animate vs. inanimate objects, using the default regularization value of .01 for covariance matrix. The decoding approach is illustrated in Figure 1B. Cross-decoding was achieved by training the animacy classifier on all trials of the intact objects experiment, and testing its discrimination of animate vs inanimate objects in the conditions of the main experiment (degraded objects). The runs of attended and unattended conditions of the intact object experiment were combined for training. Testing was performed separately for the attended and unattended conditions. Decoding was performed for every possible combination of training and testing time points between 0 and 500 ms, resulting in a 50×50 matrix of 10 ms timepoints, for each of the 4 conditions (object alone / object-in-scene x attended / unattended condition), per participant. In addition, to generate a measure of same-time cross-decoding for training and testing, decoding accuracy of each time point along the diagonal of the matrix was averaged with its neighboring time points at a radius of 2 (20 ms).

### Statistical analysis

The main hypothesis was tested using a repeated-measures ANOVA on decoding accuracies within a predefined cluster. Bayes factors for the main effects/interactions were calculated as Bayesian one-sample t-tests for the respective difference scores between conditions using the JASP default settings (Cauchy prior with scale 0.707). Above-chance decoding was also tested within the time-by-time matrices. To reduce multiple comparisons, the statistical tests in the time-by-time matrix were restricted to time points that showed significant above-chance (50%) decoding of intact objects in the intact objects experiment in a cross-validation analysis. That is, significant time points in the intact-object experiment were used to define a temporal mask for cross-decoding significance testing.

Significance was tested by computing random-effects temporal-cluster statistics corrected for multiple comparisons. This was accomplished via t-test computation over 1000 permutation iterations, in which the sign of samples was randomly flipped (over all features) after subtracting the mean, and using threshold free cluster enhancement (TFCE) as cluster statistic, with a threshold step of .1. Significance was determined at p < 0.05 by testing the actual TFCE image against the maximum TFCE scores yielded by the permutation distribution (Smith & Nichols, 2009).

## Results

### Behavioral performance

Mean accuracy in the number oddball task was 98.9% (SD=2.3) while mean accuracy in the fixation dimming task was 89.8 % (SD=10.7). The difference between tasks was significant (t25 = 4.65, p < 0.001). The overall high accuracy confirms that participants complied with task instructions. Furthermore, they show that, as expected, detecting the appearance of a number was very easy, ensuring that participants fixated the screen and attended the images (Brandman & Peelen, 2017). By contrast, the fixation dimming task was more challenging. Critically, because the fixation dims always occurred just before the image (Figure 1A), attention was temporally directed away from the image.

### Independent effects of scene context and attention on animacy cross-decoding

In our first and main analysis, we tested whether the effect of scene context observed in our previous study depended on attention. To this end, we compared the decoding accuracies within a predefined temporal cluster that previously showed contextual facilitation (Brandman & Peelen, 2017) and asked whether contextual facilitation, i.e., higher decoding accuracies for degraded objects in scenes than for degraded objects alone, would interact with attention. Object decoding accuracy was averaged across the time points of interest, at a training time of 270 ms and testing time of 320 ms, across a radius of 30 ms, as illustrated in Figure 2 (see also Brandman & Peelen, 2017). The highest decoding accuracy (57%) was observed for the attended object-in-scene condition, numerically equivalent to the decoding accuracy observed in the same condition in our previous study (Brandman & Peelen, 2017). We then tested the means in a repeated-measures ANOVA, with attention and scene context as within-subject factors. This analysis revealed main effects of scene context (F(1, 25) = 5.88, p = 0.023, η^2^ = 0.104, 95% CI [0.004, 0.055]; BF10 = 2.37) and attention F(1, 25) = 7.31, p = 0.012, η^2^ = 0.061, 95% CI [0.005, 0.040]; BF10 = 4.00), but no interaction between them (F(1, 25) = 0.73, p = 0.40, η^2^ = 0.005; BF01 = 3.46). These results replicate our previous finding of contextual facilitation and show that contextual facilitation and attention independently modulate object representations.

**Figure 2.**
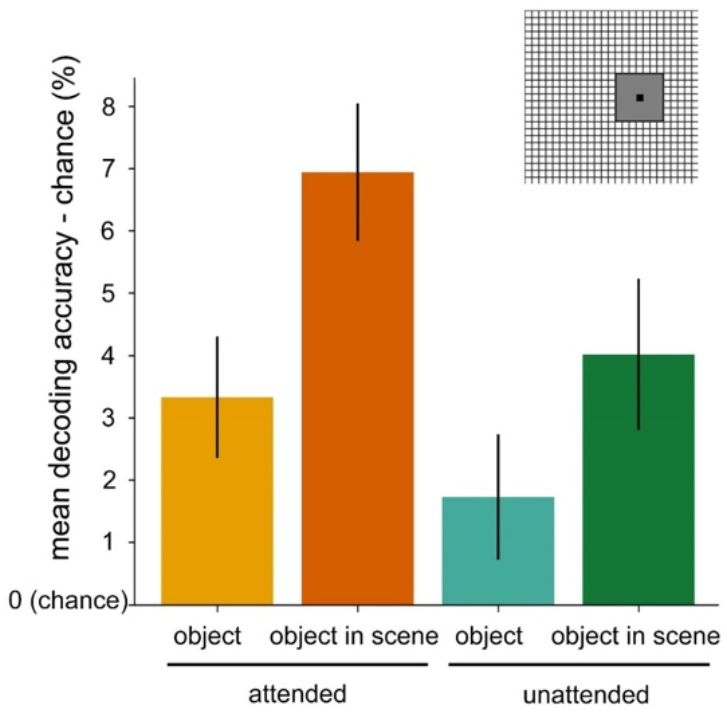
Effects of attention and scene context on object animacy decoding. Cross-decoding accuracy averaged around the previously reported peak of contextual facilitation (Brandman & Peelen, 2017; time cluster from time-by-time matrix illustrated on the top right). Results are presented as mean decoding accuracies minus chance (50% accuracy). Attention and scene context each increased object decoding accuracy independently of one another. Error bars denote the SEM across subjects.

### Cross-decoding of degraded objects

In additional exploratory analyses, we examined the time course of decoding in each condition separately, again using the cross-decoding approach (Figure 1B). In the attended condition, degraded objects alone were decoded above chance at a few isolated time points between 290 and 390 ms (Figure 3A). Degraded objects in scenes were reliably decoded across a much larger cluster of time points between 230 and 480 ms after stimulus onset (Figure 3B). In the unattended condition, there were no time points of above-chance decoding for the degraded objects alone condition (Figure 3C), while decoding for the degraded objects in scenes was above chance across a large cluster between 230 and 470 ms after stimulus onset (Figure 3D). These results again show that attention and scene context both increased decoding accuracy. However, statistical comparisons between these matrices did not yield significant time points after correcting for multiple comparisons.

**Figure 3.**
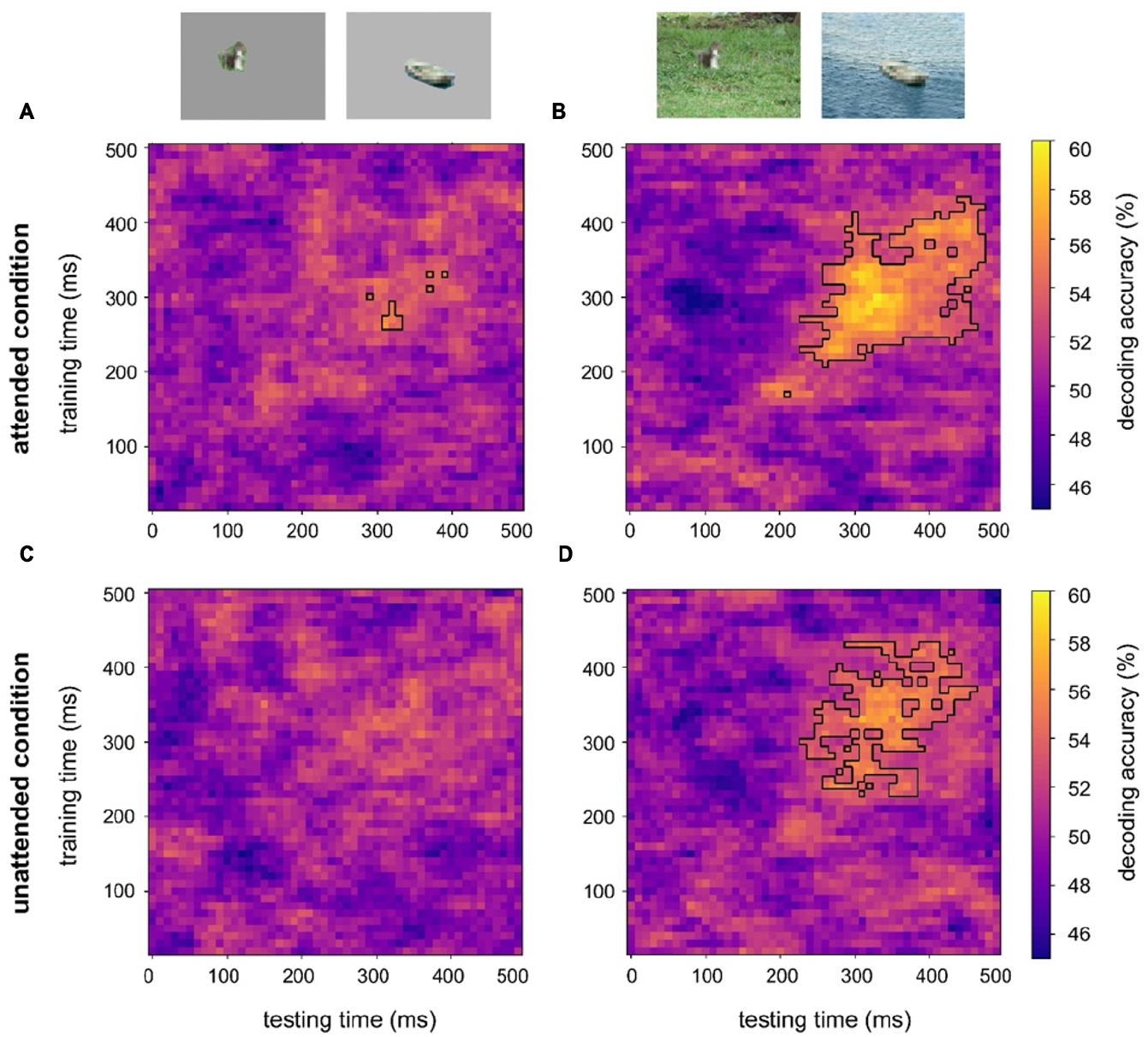
Cross-decoding of object animacy for each of the four conditions. Each time-by-time matrix presents the accuracies (mean across 26 subjects) of the animacy classifier trained on intact objects and tested on degraded objects from stimulus onset to 500 ms, in each of the four experimental conditions: **A**. Degraded objects alone in the attended condition; **B**. Degraded objects in scenes in the attended condition; **C**. Degraded objects alone in the unattended condition; **D**. Degraded objects in scenes in the unattended condition. Animacy of degraded objects alone (A, C) was decoded above chance in a small number of time points in the attended condition (A) but could not be decoded above chance in the unattended condition (C). By contrast, decoding of degraded objects in scenes (B, D) was above chance across a large cluster between 230-470ms after stimulus onset in both attended (B) and unattended (D) conditions. Black outlines represent above chance decoding (against chance, 50%; 1-tail TFCE, p<0.05).

### Contextual facilitation in a large sample of participants (N=51)

The present study made use of the same degraded-object stimuli as in our previous study (Brandman & Peelen, 2017). The attended condition was also the same as in that study. We therefore had the opportunity to concatenate the two datasets, thereby nearly doubling the number of participants (N=51) and thus attaining sufficient statistical power to test for contextual facilitation across the whole time x time matrix. The cross-decoding accuracies in the concatenated datasets are presented in Figure 4. Significant contextual facilitation (i.e., better cross-decoding of degraded objects in scenes than degraded objects alone) was found in multiple temporal clusters between 280-460 ms testing time and 220-390 ms training time after stimulus onset (Figure 4B). The contextual facilitation effect was also observed in the diagonal of the time x time matrix, with multiple significant time points between 290 and 360 ms after stimulus onset (Figure 4C). These findings provide evidence for contextual facilitation of object processing in a large sample and show that this cluster includes earlier time points (from 290 ms testing time) than previously reported (from 320 ms testing time; Brandman & Peelen, 2017).

**Figure 4.**
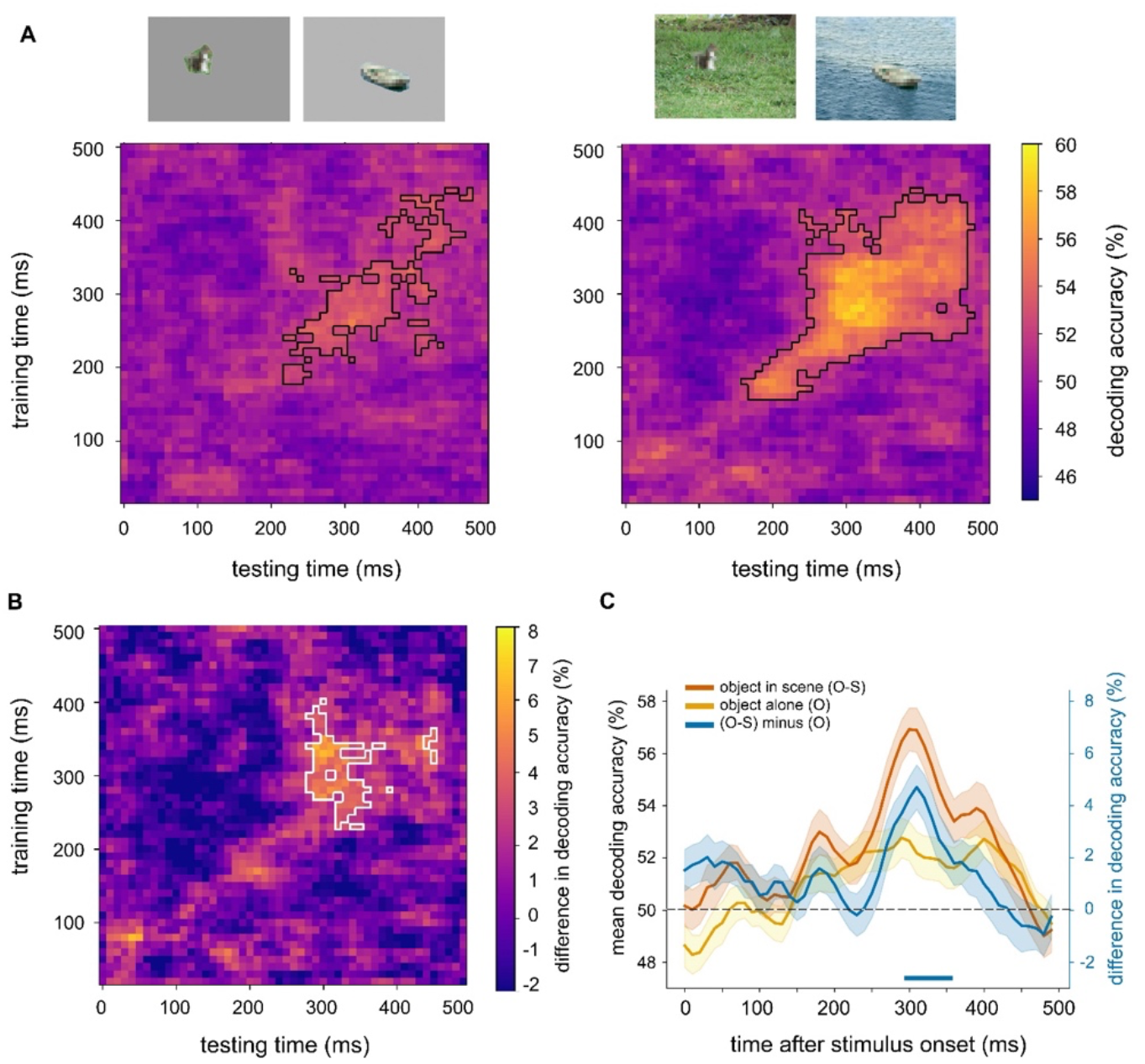
Cross-decoding of object animacy across previous and current datasets (N=51). The data from the attended condition of the present study was collapsed with the MEG dataset acquired by Brandman & Peelen (2017). A. Cross-decoding of degraded objects alone and degraded objects in scenes in the collapsed dataset (N=51). Black outlines denote above-chance decoding (TFCE against chance, p<0.05); B. Time-by-time matrix of the difference in decoding accuracies of degraded objects in scenes minus degraded objects alone. White outline denotes significantly better decoding of degraded objects in scenes than of degraded objects alone (pairwise TFCE, p<0.05); C. Cross-decoding accuracies along similar training and testing times (smoothed diagonal of matrix), for degraded objects alone (yellow), degraded objects in scenes (red), and the difference between them (blue). Decoding was significantly better for degraded objects in scenes than for degraded objects alone between 290-360 ms after stimulus onset. Dashed line marks chance-level decoding (50% accuracy). Error bars denote the SEM across subjects. Lower blue line marks the time points of significant difference between decoding of degraded objects in scenes and degraded objects alone (pairwise TFCE; p<0.05).

## Discussion

The current results provide evidence that scene context and attention independently facilitate the representation of ambiguous objects. Replicating a previous study (Brandman & Peelen, 2017), classifiers trained to distinguish clearly visible animate vs inanimate objects (without scene context) generalized to distinguish degraded objects in scenes better than degraded objects alone, despite the added clutter of the scene background. We found that attention also modulated object representations: when participants attended the scene, decoding of object category was better than when participants attended the fixation cross. Notably, we found substantial evidence (BF>3) that these two effects were independent of each other. Finally, combining the data of the two studies (N=51) allowed for investigating the time course of context-based facilitation with high statistical power, revealing a large cluster of significant facilitation that started at around 220 ms training time and 280 ms testing time.

We interpret these findings as evidence that expectations derived from scene context feed back to modulate object representations, such that the representation of an ambiguous (e.g., blurred) object, at around 300 ms after scene onset, becomes more similar to the representation of its intact (i.e., sharp) version slightly earlier in time. This interpretation is in line with the behavioral finding that blurred objects in scenes are perceived as sharper than the same objects outside of scene context (Rossel et al., 2022, 2023). The time course revealed here, with facilitation starting at 280 ms (in the time x time matrix analysis) or 290 ms (in the diagonal analysis), matches the time course of a previous transcranial magnetic stimulation (TMS) study, which showed that stimulation of object-selective cortex at 280-300 ms after stimulus onset impaired the recognition of degraded objects in scenes (Wischnewski & Peelen, 2021).

In the unattended condition of the current study, the scenes were task-irrelevant, with participants monitoring the dimming of the fixation cross instead. Furthermore, this dimming always occurred just before scene onset, such that participants had to also temporally direct their attention away from the scene. The finding that this attention manipulation did not interact with the context manipulation is in line with previous behavioral and EEG studies that showed that semantic congruency effects can occur independently of attention and task relevance (Chen et al., 2022; Cornelissen & Võ, 2017; Munneke et al., 2013). However, unlike these previous studies, here we investigated how the representations of ambiguous objects were modulated by scene context, rather than whether semantic violations of clearly-visible objects were automatically noticed. As reviewed in the Introduction, these two processes may rely on different neural mechanisms.

While the current attention manipulation was effective, as demonstrated by its effect on decoding accuracy, we cannot rule out that stronger manipulations of attention would additionally reveal an interaction between context and attention. For example, the within-subject design may have led to some spill-over between the tasks, such that participants may have continued to pay attention to the scenes in the unattended condition blocks (note, however, that the fixation dimming task was quite challenging, arguing against this possibility). Furthermore, the objects in the scenes were presented close to fixation, such that spatial attention was likely directed to them. Finally, the scenes only contained one foreground object, resulting in a low perceptual load. Previous studies have shown that increasing attentional and/or perceptual load results in attentional selection at increasingly early stages of visual processing (Lavie, 2005). Accordingly, it is possible that context-based facilitation would be reduced when attentional selection occurs at an earlier processing stage due to high attentional or perceptual load. Nevertheless, our results show that the representation of an object is facilitated by scene context even when the object is entirely irrelevant to the task at hand.

Our results appear to contradict those of a recent EEG study that compared EEG responses evoked by scene-consistent objects (e.g., a computer monitor on a desk) with responses evoked by scene-inconsistent objects (e.g., a computer monitor on a kitchen counter; Chen et al., 2022). In that study, the decoding of object category was enhanced for semantically consistent (vs. inconsistent) objects only when the objects were task relevant. By contrast, in the current study, we found that context-based facilitation was independent of task relevance. An important difference between these studies is that the objects in the task-irrelevant condition of Chen et al. (2022) were not degraded as in the current study and were fixated for a relatively long time (1500 ms vs. 50 ms in the current study). The lack of a difference in the decoding of semantically consistent versus inconsistent objects in that study may therefore reflect the fact that there were no features to be disambiguated by scene context. Furthermore, in the Chen et al. study, the task-irrelevant condition differed from the task-relevant condition in important aspects, with the objects presented for 1500 ms in the task-irrelevant condition but only very briefly (83 ms) in the task-relevant condition. This raises the possibility that the effect of scene context on object decoding depends on presentation time rather than task relevance. More generally, context may facilitate decoding only when bottom-up object information is unreliable, for example because of brief presentation time or because of degradation. This notion is consistent with previous behavioral findings, showing smaller facilitation effects of scene context for intact than for degraded object detection (Brandman & Peelen, 2017). Taken together, we propose that context-based facilitation of object processing is stimulus dependent rather than task dependent.

Which could be the scene properties that drive context-based facilitation? In the current study, there were multiple scene properties that may have contributed to the facilitation observed. For example, the category of the scene (e.g., a lake) and the position of the object in the scene (e.g., in the sky) both provide information about the likely category of the ambiguous object (Biederman et al., 1982; Davenport & Potter, 2004). Furthermore, the approximate distance of the object in the scene provides information about its real-world size, which has been shown to inform the recognition of degraded objects (Gayet et al., 2022). While we did not manipulate these scene properties here, the current finding that context modulated object representations independently of attention indicates that the properties resulting in facilitation were extracted quickly and automatically. This is in line with previous studies showing that many scene properties (e.g., scene category) can be extracted with limited focused attention (Evans et al., 2011; Li et al., 2002; Potter, 1976). Future studies are needed to systematically manipulate informative scene properties and test which of these drive the automatic facilitation observed here.

More generally, the current results speak to the relation between attention and expectation in the context of naturalistic vision. Previous studies have shown that attention and expectation can both facilitate perception (Summerfield & Egner, 2009), raising the possibility that the previously observed facilitatory effects of scene context (Brandman & Peelen, 2017) reflected an effect of attention rather than expectation. For example, instructing participants to look for a specific object or object category in a scene enhances the representation of these objects in visual cortex from around 180 ms after scene onset (Kaiser et al., 2016), reflecting an influence of top-down attention (Peelen & Kastner, 2014). The current finding that the facilitatory effects of contextual expectations on object representations were independent of attention makes it unlikely that these effects were mediated by increased attention to objects, i.e., that participants voluntarily directed attention to objects that were expected in a particular context. At a neural level, the independence of the effects of attention and expectation shown here implies that contextual feedback and attentional feedback to visual cortex are mediated by different pathways. One possibility is that scene representations in scene-selective visual cortex (including the parahippocampal cortex; Aminoff et al., 2013; Bar, 2004) interact with object representations in object-selective visual cortex without involving the frontal-parietal network associated with top-down attention (Corbetta & Shulman, 2002). Contextual expectations may additionally involve the medial prefrontal cortex (Aminoff et al., 2013; Bar, 2004).

The finding that the effects of attention and expectation were additive rather than interactive is in line with previous M/EEG decoding studies using simplified stimuli (Alilović et al., 2019; Moerel et al., 2022; Smout et al., 2019). For example, in one study, feature-based attention enhanced decoding of stimulus orientation from 230 ms after stimulus onset, an effect that was independent of temporal expectations (Moerel et al., 2022). Another study compared the representation of stimulus orientation between gratings for which orientation was unexpected (i.e., surprising) with gratings for which orientation was unpredictable (Smout et al., 2019). Attention was manipulated similarly to the current study, by making the gratings task-relevant (detecting target gratings with a higher spatial frequency) or -irrelevant (detecting fixation dims). Attention similarly enhanced orientation representations of both surprising and unpredictable gratings, providing further evidence for a dissociation between attention and expectation. Nevertheless, while these and other studies showed that attention and expectation can be dissociated, there are also several ways in which attention and expectation interact. For example, attention increases the representation of unexplained (or mismatching) perceptual input (prediction error; (Jiang et al., 2013; Smout et al., 2019) and, conversely, unexpected stimuli (e.g., a semantically inconsistent object in a scene) can attract attention (Spaak et al., 2022; Underwood & Foulsham, 2006). Attention can also be guided by contextual expectations about object location or appearance, for example during visual search in scenes (Castelhano & Krzyś, 2020; Gayet & Peelen, 2022; Peelen & Kastner, 2014; Võ et al., 2019). These studies point to an intricate relation between attention and expectation that requires further study.

To conclude, our findings suggest that contextual expectations that have been learned over many years affect object processing relatively automatically, i.e., independently of attention and task relevance. Taken together with previous findings (Peelen et al., 2024), our results indicate that facilitatory scene-object interactions depend on the (un)reliability of the visual input more than on the specific task performed on this input.

## Acknowledgements

This project received funding under the European Union’s Horizon 2020 research and innovation program, from the European Research Council (ERC) (grant agreement No. 725970) and the Marie Sklodowska-Curie (grant agreement no 659778).

